# Tumor microenvironment exploration and therapeutic target identification in bladder cancer based on Cuproptosis signature and chemotherapy-resistant model: results from bioinformatics and experimental validation

**DOI:** 10.1101/2023.05.23.542015

**Authors:** Jun Gu, Zhen-duo Shi, Kun Pang, Lin Hao, Wei Wang, Cong-hui Han

## Abstract

**Background:** The discovery of cuproptosis provides a new way to make full use of the pathophysiological effects of copper for anticancer therapy and could help identify a therapeutic target in bladder cancer.

**Methods:** In this study, 411 BLCA tumor samples from the Cancer Genome Atlas (TCGA) cohort were obtained. Differentially expressed genes (DEGs) between chemotherapy-sensitive and chemotherapy-resistant mouse bladder cancers were also obtained. Sixteen genes were defined as cuproptosis-related genes (CRGs), and the cutoff score was calculated based on LASSO Cox regression. CCK-8 and Transwell assays were used to detect the migration and proliferation of 5637 and T24 cells, respectively. Liarozole dihydrochloride (L-D), a mild P450, inhibits the expression of CYP26B1. Multiple immunohistochemistry analyses were used to explore the association between the immune microenvironment and CRGs.

**Results:** A higher Cuproptosis score was significantly associated with worse overall survival in BLCA (**p**<0.0001, HR=2.32). Single-cell transcriptional data were used to assess the function of *CYP26B1* and *CYP26B1* may be negatively associated with DNA damage and repair. In vitro experiments indicated that overexpression of CYP26B1 enhanced the migration and proliferation of BLCA cells, while L-D inhibited the migration and proliferation of BLCA cells. A higher expression level of Dihydrolipoamide S-Succinyltransferase (*DLST*) serves could as a risk factor in patients treated with atezolizumab and higher expression of *DLST* suggest an immunosuppressive microenvironment.

**Conclusions:** *CYP26B1* may be a therapeutic target for bladder cancer, and the higher expression of DLST may suggest an immunosuppressive microenvironment in BLCA.

**Key Messages:** Cuproptosis may be associated with the chemotherapy-resistance of bladder cancer and our findings may provide new ideas for the treatment of bladder cancer.

## Introduction

Ferroptosis, first described by Dixon et al., is a unique cell death pathway driven by iron-dependent lipid peroxidation [1]. Owing to the high iron levels in cancer cells and increased sensitivity to ferroptosis induction, inducing ferroptosis is generally considered a promising cancer treatment[2, 3]. Recently, Tsvetkov et al. first proposed a new cell death pathway, “cuproptosis,” based on the important role of copper ions in the human body [4]. Copper ions directly bind to the fatty acylated components of the tricarboxylic acid (TCA) cycle, resulting in abnormal aggregation of fatty acylated proteins and loss of iron-sulfur proteins, which further leads to protein toxic stress reactions and eventually cell death [4]. Because the role of ferroptosis in various cancers has been widely studied, it may also play a role in tumor occurrence and development. The discovery of cuproptosis provides a new method for fully exploiting the pathophysiological effects of copper in anticancer therapy.

Bladder cancer (BLCA) is one of the ten most prevalent cancers worldwide and the second-largest urinary system tumor [5, 6], of which urothelial non-muscle invasive bladder cancer (NMIBC) is the most common. Patients with NMIBC have a high risk of recurrence (approximately 50%–70%), followed by progression to muscle-invasive bladder cancer (MIBC) [7, 8].Therefore, most patients undergo lifelong cystoscopic monitoring and various treatment interventions, making BLCA treatment one of the most expensive. Systemic chemotherapy is used to control BLCA progression and alleviate its symptoms. Cisplatin-based chemotherapy is the first-line treatment for most cancers. Although this treatment can theoretically reduce the risk of death from BLCA in clinical practice, the overall efficacy rate is less than 50% [9, 10]. For a long time, little progress has been made in BLCA treatment [10]. In recent years, new treatment options, such as PD-1/PD-L1 inhibitors, have emerged. However, immunotherapy for BLCA remains immature, and more efforts are urgently needed [11]. Therefore, the development of new strategies for BLCA treatment is of great clinical significance.

Bioinformatics provides a vast and convenient platform for studying the pathogenesis of various diseases, especially cancer, using large databases, powerful analytical tools, and evaluation methods [12].In this study, we explored the function of cuproptosis-related genes (CRGs) in chemotherapy resistance and the immune microenvironment of BLCA, using various public datasets and a series of experiments to identify pivotal CRGs that can effectively predict chemotherapy resistance and immune infiltration in BLCA. Our findings may contribute to the development of novel and precise therapies for BLCA and provide new ideas for its treatment.

## Materials and methods

### Data collection and normalization

In this study, gene expression profiles and corresponding clinical information of 411 BLCA tumor samples were obtained from the Cancer Genome Atlas (TCGA) cohort (https://portal.gdc.cancer.gov/). Wang et al. [13] established chemotherapy-sensitive and chemotherapy-resistant MIBC models. We obtained differentially expressed genes (DEGs) between chemotherapy-sensitive and chemotherapy-resistant mouse BLCA and the GSE192575 dataset from the Gene Expression Omnibus (GEO) database. We also obtained the IMvigor210 cohort from http://research-pub.gene.com/IMvigor210CoreBiologies, which included patients with locally advanced and metastatic uroepithelial cancer treated with atezolizumab. Single-cell RNA-seq profiling of two BLCA specimens was performed using the GSE130001 dataset (repeated samples and samples with a follow-up time of fewer than 30 d were deleted).

### Exploring the potential function of CRGs in BLCA

Sixteen genes have been identified as CRGs: *FDX1*, *LIAS*, *LIPT1*, *DLD*, *DLAT*, *PDHA1*, *PDHB*, *MTF1*, *GLS*, *CDKN2A*, *DBT*, *GCSH*, *DLST*, *SLC31A1*, *ATP7A*, and *ATP7B* [4].

The cBioPortal for Cancer Genomics (http://cbioportal.org) provides a web resource for exploring, visualizing, and analyzing multidimensional cancer genomic data [14]. We used cBioPortal to display the deletions, amplifications, and other genomic variations in the CRGs of BLCA. Gene set cancer analysis (GSCA) is an integrated platform for genomic, pharmacogenomic, and immunogenomic GSCA [15]. In this study, we used the GSCA tool to explore the expression levels and potential prognostic significance of CRGs in BLCA. Additionally, we used the GeneMANIA prediction server [16] (biological network integration for gene prioritization and gene function prediction) to construct an interaction network and determine the potential biological functions of the CRGs.

### Calculating the cuproptosis score and identifying potential therapeutic target

We used LASSO Cox regression to screen and explore the most important prognostic CRGs and calculated the cutoff score based on these results. Based on the median cutoff value, we compared the overall survival between the low and high cutoff groups. In addition, we used the LIMMA package [17] to identify the DEGs between the low and high cutoff score groups. Genes with logFC ≥ 1 and p-value < 0.05 were considered significant DEGs. We used WebGestalt [18] (WEB-based GEne SeT AnaLysis Toolkit, http://www.webgestalt.org/), a functional enrichment analysis web tool, to determine the potential biological functions of the DEGs between the two groups. We further used the DEGs between chemotherapy-sensitive and chemotherapy-resistant mouse bladder cancer (GSE192575) to identify common DEGs in the high cutoff score group and chemotherapy-resistant BLCA. We also used the STRING tool [19] (STRING v11: protein–protein association networks with increased coverage, supporting functional discovery in genome-wide experimental datasets) to construct a protein–protein interaction network.

### Exploring the potential function of the therapeutic target

Cytochrome P450 proteins are monooxygenases that catalyze many reactions in drug metabolism and cholesterol, steroid, and other lipid syntheses. Among the 74 common DEGs in the high cutoff score group and chemotherapy-resistant BLCA, we found a gene (cytochrome P450 family 26 subfamily B member 1, *CYP26B1*) that encodes a member of the cytochrome P450 superfamily and explored its potential function. The CanserSEA tool [20] (a dedicated database that comprehensively decodes distinct functional states of cancer cells at a single-cell resolution) was used to determine the potential biological function of *CYP26B1* in various tumor types. The LinkedOmics (a publicly available portal that includes multi-omics data from all 32 TCGA cancer types) tool [21] was used to identify the most relevant genes with *CYP26B1*, and functional enrichment analysis was used to predict biological function.

### Cell culture and transfection

Two human BLCA cell lines (5637 and T24) were maintained in our laboratory. The cells were cultured in RPMI 1640 medium containing 10% fetal bovine serum (FBS). Cells were maintained at 5% CO2 at 37 °C. To explore the potential function of *CYP26B1* in BLCA, 5637 and T24 cells were transfected with an empty vector or *CYP26B1*-overexpression plasmid using Lipofectamine 3000 reagent (Invitrogen, Waltham, MA, USA), according to the manufacturer’s instructions. Additionally, we used liarozole dihydrochloride (L-D, Selleck Chemicals, Houston, TX, USA, number: S6594), a mild inhibitor of P450, to inhibit *CYP26B1* expression.

### CCK-8 and Transwell assays

For the CCK-8 assay, 1 × 10^3^ 5637 or T24 cells were seeded into each well of a 96- well plate. We measured the proliferation of 5637 and T24 cells from days 1 to 5 using a CCK-8 kit (Dojindo, Kumamoto, Japan). For the Transwell assay, we seeded 3 × 10^4^ 5637 or T24 cells (200 μL FBS-free medium) in the upper well of the Boyden Transwell chambers and added 800 µL of culture medium supplemented with 10% FBS to the bottom chamber. Cells were cultured for 48 h at 37 ℃, then the cells on the upper surface of the chamber were wiped off, and the cells on the lower surface were fixed with 4% paraformaldehyde solution and stained with 0.1% crystal violet.

### Exploring the associations between CRGs and the tumor microenvironment

The association between CRGs (gene set levels) and tumor-infiltrating immune cells was evaluated using the GSCA tool. The IMvigor210 cohort included patients with locally advanced and metastatic uroepithelial cancer treated with atezolizumab, and the prognostic value of CRGs in the IMvigor210 cohort was evaluated. In addition, the associations between dihydrolipoamide S-succinyltransferase (*DLST*) and immune stimulators and inhibitors were evaluated using the TISIDB [22] tool (a web portal for tumor and immune system interactions, which integrates multiple heterogeneous data types).

### External validation using multiplex immunohistochemistry (mIHC)

The BLCA tissue microarrays were obtained from Shanghai Wellbio Biotechnology Co., Ltd. (Shanghai, China). To explore the association between *DLST* expression and tumor-infiltrating immune cells (CD3^+^, CD4^+^, and CD8^+^ T cells), mIHC was performed using anti-human CD3 (Abcam, Cambridge, UK; ab135372), CD4 (Abcam; ab133616), CD8 (Abcam; ab217344), and DLST antibodies (Abcam; ab177934). The Visiopharm software (Visiopharm A/S, Hersholm, Denmark) was used to quantify the expression levels of DLST, CD3, CD4, and CD8.

### Statistical analysis

The fold change of DEGs was calculated as mean (tumor)/mean (normal), and the p-value was estimated by t-test and was further adjusted by false discovery rate (FDR). The Cox proportional hazards model and log-rank tests (Kaplan–Meier method, median mRNA value was set as the cutoff value) were performed for CRGs in BLCA. To evaluate pathway activity, samples were divided into two groups (high and low cutoff) by median gene expression, and the difference in the pathway activity score (PAS) between groups was defined using the Student’s t-test. The *p*-value was adjusted by FDR, and FDR ≤ 0.05 was considered significant.

## Results

### The potential function of CRGs in BLCA

To explore the potential functions of CRGs (*FDX1*, *LIAS*, *LIPT1*, *DLD*, *DLAT*, *PDHA1*, *PDHB*, *MTF1*, *GLS*, *CDKN2A*, *DBT*, *GCSH*, *DLST*, *SLC31A1*, *ATP7A*, and *ATP7B*) in BLCA, we first detected genomic variations in cBioPortal. The general mutation rate of the CRGs in BLCA was relatively high, and 35%, 5%, and 2.9% of BLCA samples had variants of *CDKN2A*, *ATP7B*, and *MTF1*, respectively (**Figure 1A**). Most variants of *CDKN2A* and *ATP7B* were deep deletions, whereas *MTF1* variants were mostly amplified. The relatively high mutation rate of CRGs in BLCA suggests that CRGs play an important role in BLCA development. Next, we explored the expression levels of CRGs and found that the expression levels of *SLC31A1*, *CDKN2A*, and *GCSH* were significantly higher in tumor samples, and those of *DLST* and *MTF1* were significantly lower in tumor samples (**Figure 1B**). We further explored the associations between CRGs and various biological functions and found that CRGs may be significantly associated with apoptosis and the cell cycle (*p*<0.05, **Figure 1C**). Additionally, *SLC31A1*, *PDHA1*, and *MTF1* may be positively associated with apoptosisand the cell cycle, whereas *LIAS*, *DBT*, and *ATP7A* may be negatively associated with apoptosis and EMT (**Figure 1E**). Univariate regression analysis was used to evaluate the prognostic significance of the CRGs. The expression levels of *SLC31A1* and *GCSH* were negatively associated with patient prognosis, including disease-specific survival (DSS), overall survival (OS), and progression-free survival (PFS) (**Figure 1D**). The expression levels of *LIPT1* were significantly associated with longer DSS, OS, and PFS. **Figure 1F–H** illustrates the overall survival curves, and the cutoff was selected as the median expression of *GCSH*, *SLC31A1*, and *LIPT1*.

**Figure 1.**
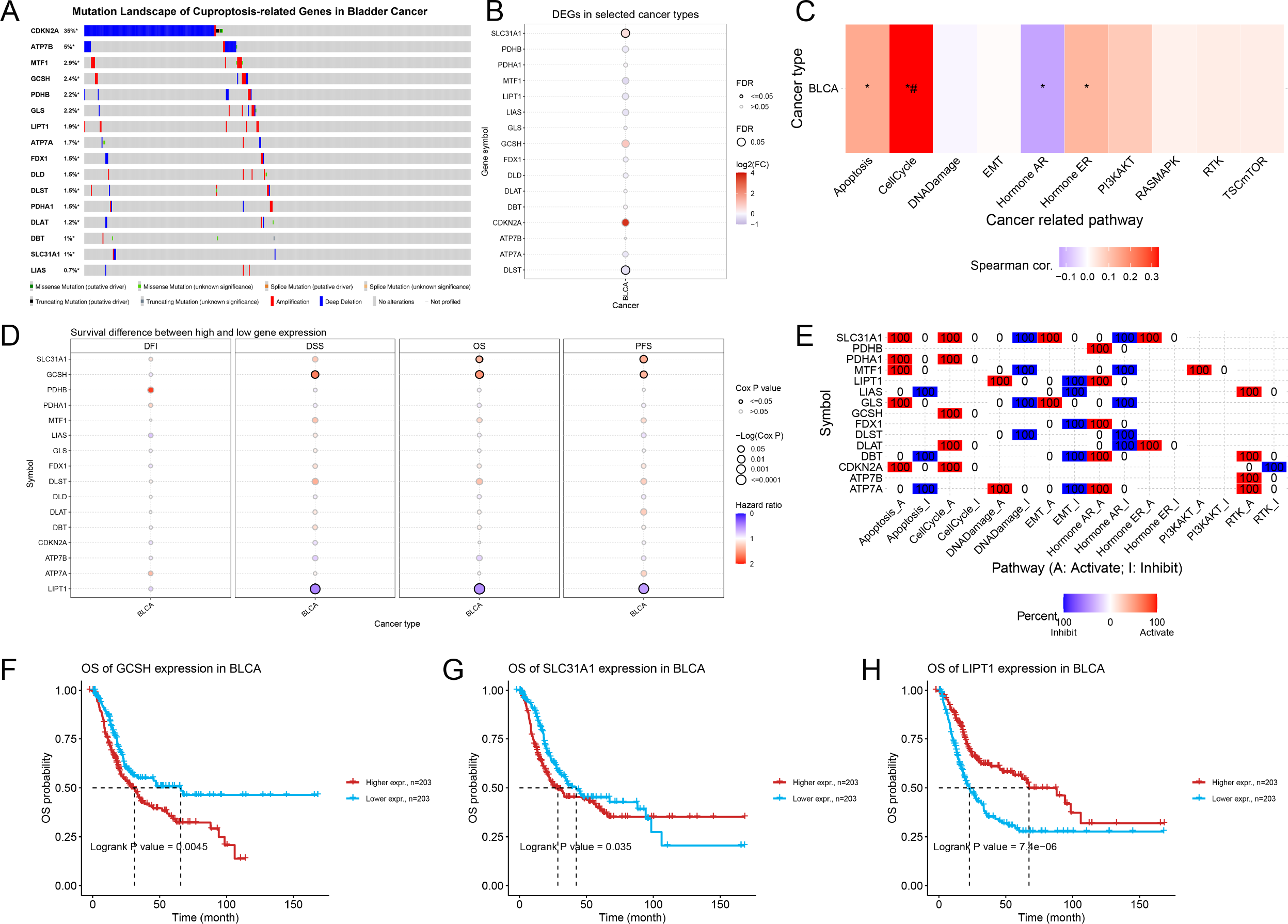
The potential function of CRGs in BLCA. (A)The genome variation of 16 CRGs in bladder patients from cBioPortal. (B) The expression level of 16 CRGs in bladder patients from TCGA datasets (p-value was estimated by t-test and was further adjusted by FDR). (C) The associations between CRGs and various biological function in cancer (student T test, P value was adjusted by FDR). (D) Evaluation of the prognostic significance of CRG using univariate regression analysis (Cox Proportional-Hazards model and Log-rank tests). (E) The potential biological functions in tumors corresponding to each CRG. (F-H) Overall survival curves of GCSH, SLC31A1 and LIPT1 in bladder cancer, respectively (Log-rank tests).

### Construction of the interaction network and calculating the cuproptosis score

The CRGs had a close and complex interaction (including physical interaction, shared protein domains, and co-expression) and function in the tricarboxylic acid cycle enzyme complex, oxidoreductase complex, and so on (**Figure 2A**). To further screen and construct a simple score to assess apoptosis, we used LASSO Cox regression to establish a cuproptosis cutoff score (**Figure 2B–C**). Based on the median value of the cutoff score, samples from the TCGA-BLCA cohort were divided into low- and high- proptosis score groups. Higher cuproptosis scores were significantly associated with worse OS in BLCA (*p*<0.0001, HR=2.32, **Figure 2D**). Furthermore, DEGs between the low and high cuproptosis score groups were identified, and functional enrichment analysis indicated (**Figure 2E)** that a high cutoff score is associated with tissue development (GO:0009888), epithelium development (GO:0060429), and epithelial cell differentiation (GO:0030855).

**Figure 2.**
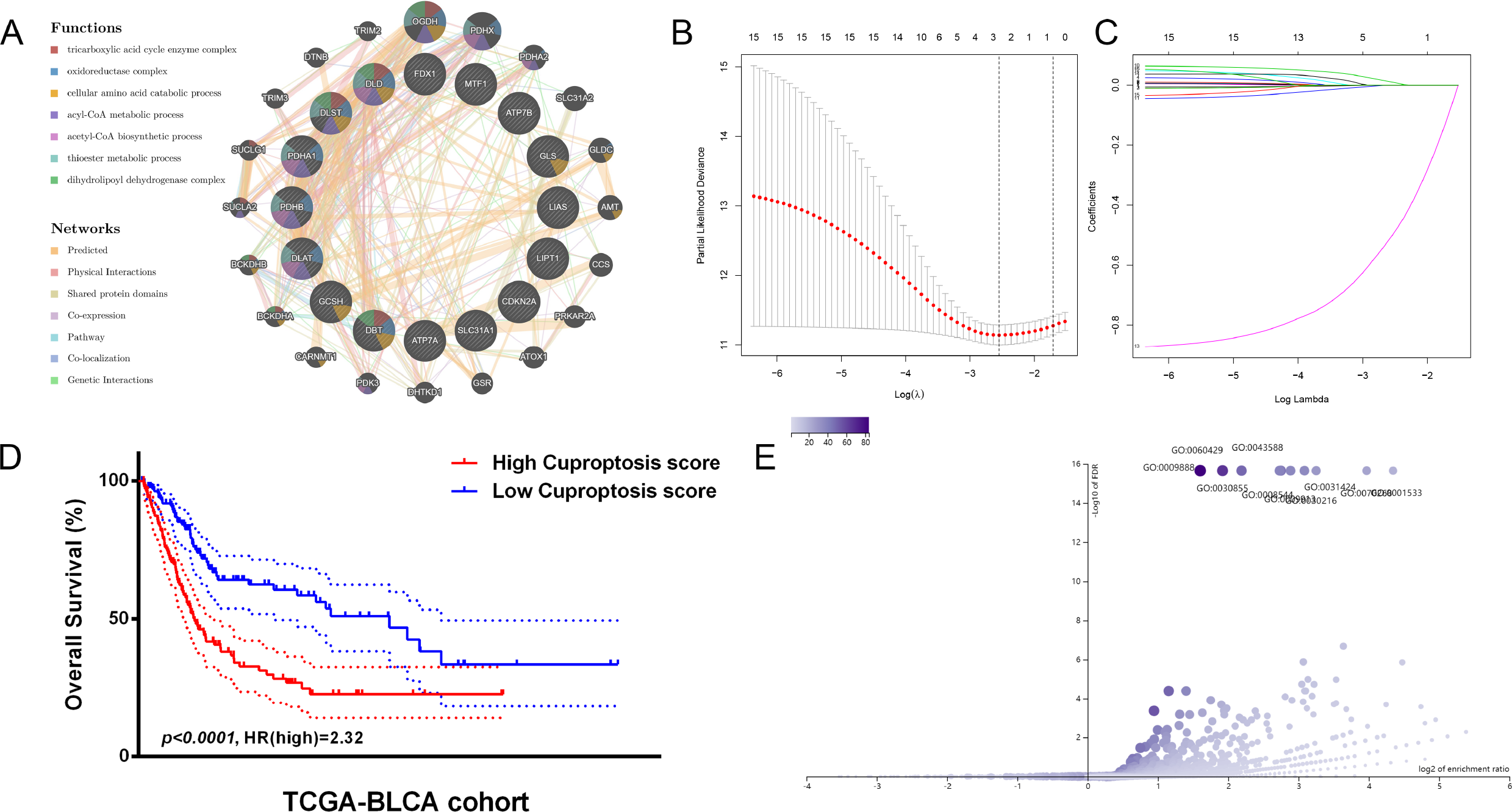
Construction of interaction network and Calculating the cuproptosis score. (A) Interaction between CRGs using the GeneMANIA tool. (B and C) Establishing cuproptosis scores using lasso cox regression. (D) Overall survival curves for the high and low cuproptosis score groups. (***p***<0.0001, HR=2.32, Log-rank tests) (E) Differential genes between high and low cuproptosis score groups and their functional enrichment results.

### Exploring potential therapeutic target associated with cuproptosis and chemotherapy-resistant BLCA

We enrolled a cohort (GSE192575) containing chemotherapy-resistant and chemotherapy-sensitive BLCA models to explore the therapeutic targets associated with drug resistance. The DEGs between the high and low cuproptosis score groups are presented in **Figure 3A**, whereas those between chemotherapy-resistant and chemotherapy-sensitive BLCA models are presented in **Figure 3B**. The Venn diagram was used to identify 74 common DEGs (**Figure 3C**). Next, we constructed a protein– protein interaction network and found that these common DEGs may still be associated with the tricarboxylic acid cycle enzyme complex and oxidoreductase complex (**Figure 3D–E**). Furthermore, functional enrichment analysis indicated that these common DEGs play important roles in the ErbB signaling pathway, estrogen signaling pathway, and cell adhesion (**Figure F–G**).

**Figure 3.**
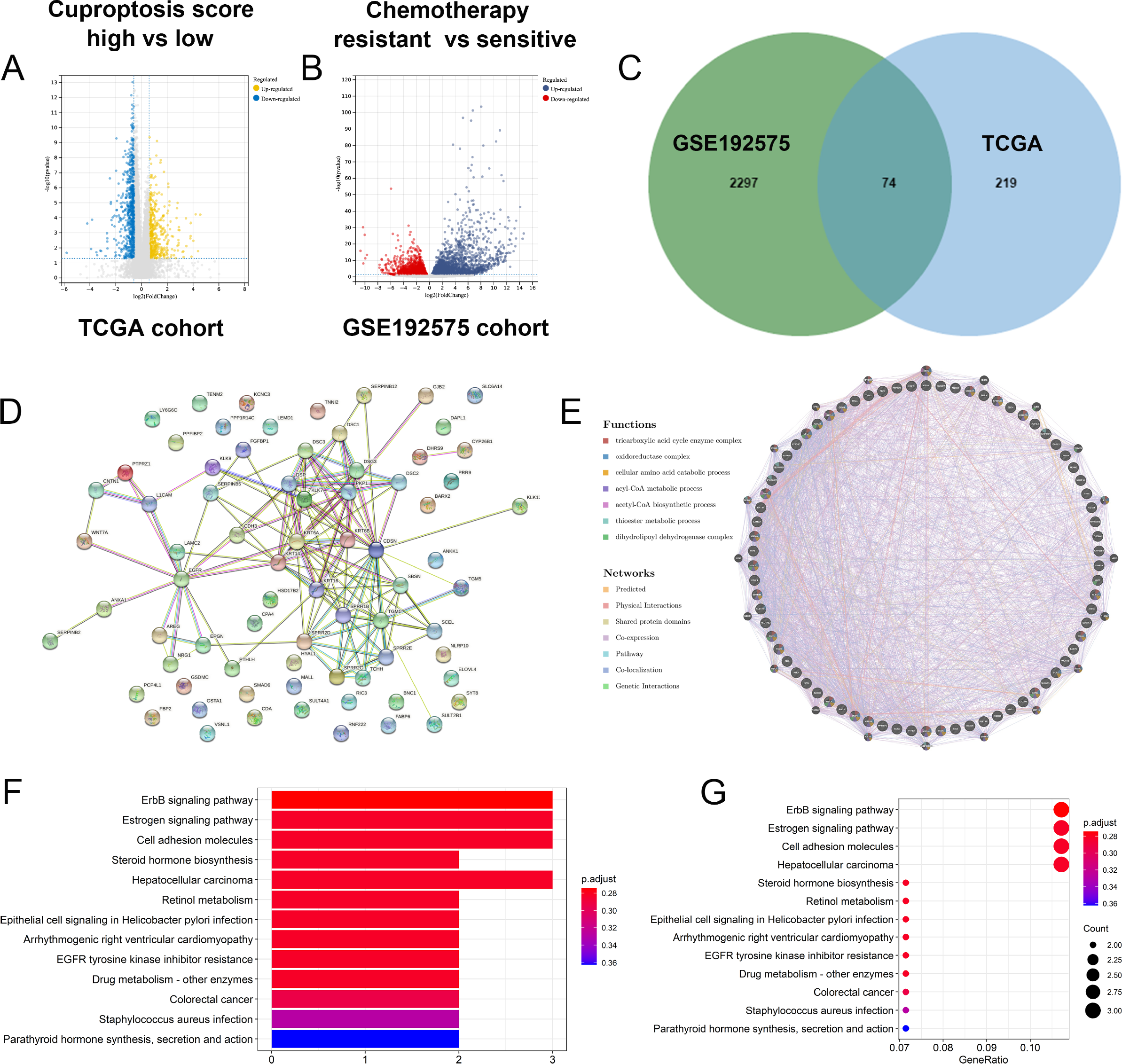
Explore potential therapeutic target associated with Cuproptosis and chemotherapy-resistant. (A) The DEGs between high and low Cuproptosis score group. (B) DEGs between chemotherapy-resistant and sensitive BLCA. (C) Venn diagrams showing the DEGs common in (A) and (B). (D and E) Protein interaction network diagram constructed based on the above DEGs. (F and G) Go function enrichment of the above DEGs and KEGG pathway enrichment results.

### Identifying CYP26B1 as a potential therapeutic target

Among the 74 common DEGs in the high cutoff score group and chemotherapy-resistant BLCA, *CYP26B1* may be a therapeutic target for BLCA. Single-cell transcriptional data were used to assess the function of *CYP26B1*, which indicated that *CYP26B1* is negatively associated with DNA damage and repair (**Figure 4A**). As DNA repair plays a key role in malignant tumor development, *CYP26B1* may promote BLCA development. An external cohort was used to explore the prognostic value of *CYP26B1*. Higher expression levels of *CYP26B1* were significantly associated with worse OS (*p*<0.01, HR=3.414, **Figure 4B**) and DSS (*p*<0.01, HR=3.08, **Figure 4C**). Further analysis (**Figure 4D–E**) indicated that *CYP26B1* was significantly negatively associated with DNA repair (correlation=–0.58, *p*<0.001), DNA damage (correlation=– 0.51, *p*<0.001), and apoptosis (correlation=–0.42, *p*<0.001). As homologous recombination deficiency (HRD) and microsatellite instability (MSI) may be associated with DNA repair, we evaluated the correlation between *CYP26B1* and HRD and MSI scores. *CYP26B1* was positively associated with HRD scores in BLCA and breast cancer (BRCA) (**Figure 4F**). *CYP26B1* (**Figure 4G**) may be positively associated with MSI scores in BRCA and colon adenocarcinoma (COAD). Thus, a higher expression level of *CYP26B1* may indicate an unstable genomic state and cause malignant tumors.

**Figure 4.**
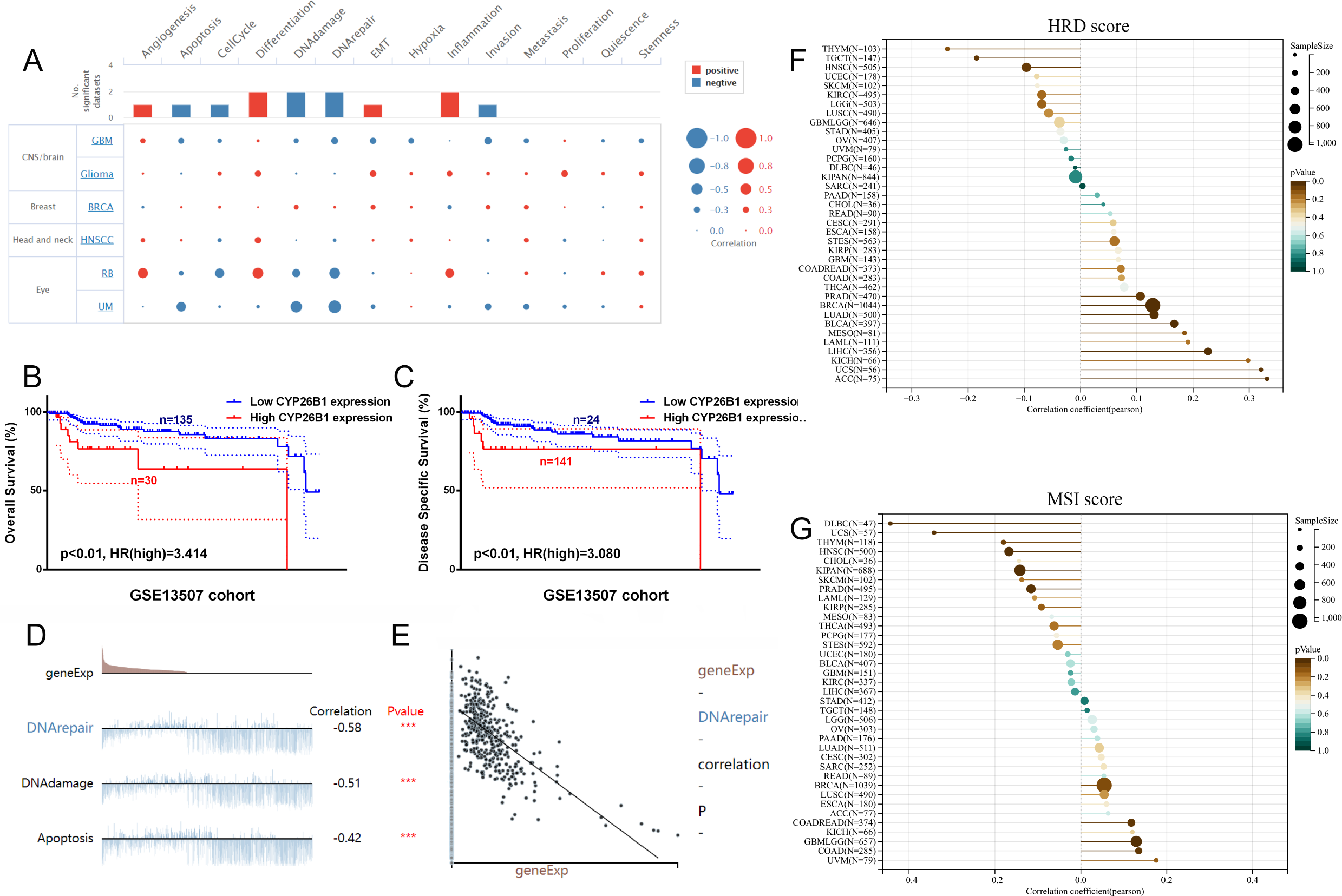
Identifying *CYP26B1* as a potential therapeutic target. (A) Single-cell transcriptional data to assess the potential function of CYP26B1 in bladder cancer. (B and C) External cohort validation of OS and DSS of cytochrome P26B1 in bladder cancer, respectively (Log-rank tests). (D and E) The correlation between expression level of CYP26B1 and DNA repair (Spearman correlation test). (F) The correlation between CYP26B1 and HRD score (Spearman correlation test). (G) the correlation between *CYP26B1* and MSI score (Spearman correlation test).

### CYP26B1 promotes BLCA migration and proliferation in 5637 and RT4 cells

In vitro experiments were performed to explore the potential function of *CYP26B1* in BLCA using MIBC (5637) and NMIBC (RT4) cell lines. Western blotting indicated that the CYP26B1 inhibitor (L-D) significantly decreased the expression level of *CYP26B1* in both 5637 (**Figure 5A**) and RT4 cell lines (**Figure 6A**) and confirmed the overexpression efficiency. Transwell and wound healing assays indicated that *CYP26B1* overexpression enhanced the migration of 5637 and RT4 cells, whereas L-D inhibited the migration of 5637 cells (**Figure 5B–C**) and RT4 cells (**Figure 6B–C**). CCK-8 and colony formation assays indicated that *CYP26B1* overexpression enhanced the proliferation of 5637 cells and RT4 cells, whereas L-D inhibited the proliferation of 5637 cells (**Figure 5D–F**) and RT4 cells (**Figure 6D–F**). These results indicate that *CYP26B1* enhances BLCA migration and proliferation.

**Figure 5.**
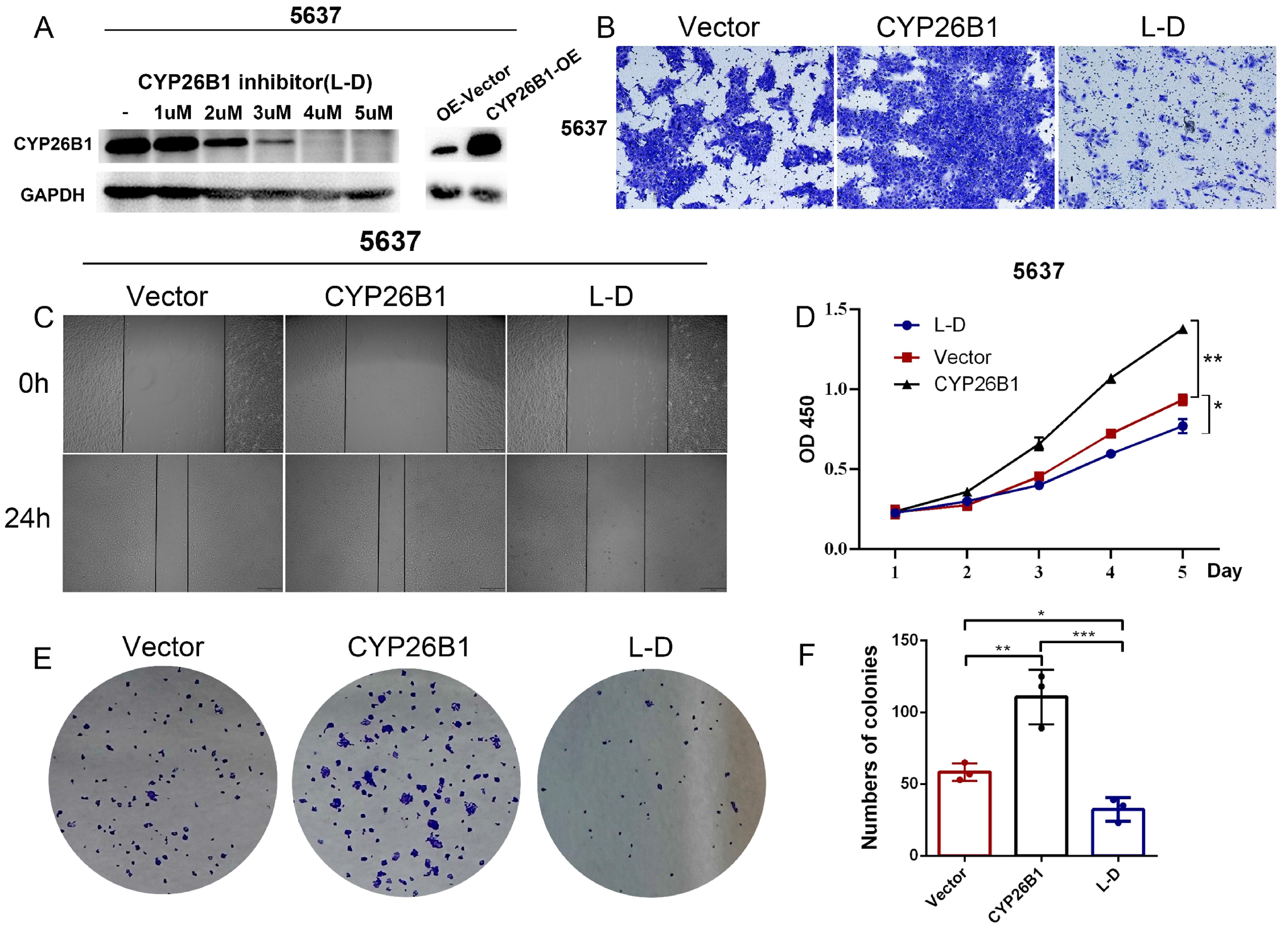
CYP26B1 promote BLCA migration and proliferation in 5637 cells. (A) Validation of inhibition and overexpression efficiency by western blot. (B) Transwell assay to explore cell migration ability in 5637 cell after overexpression and inhibition of CYP26B1. (C) Wound-healing assay to explore cell migration ability in 5637 cell after overexpression and inhibition of CYP26B1. (D) CCK8 assay to explore proliferation ability in 5637 cell after overexpression and inhibition of CYP26B1. (E-F) Colony formation assay to explore proliferation ability in 5637 cell after overexpression and inhibition of CYP26B1.

**Figure 6.**
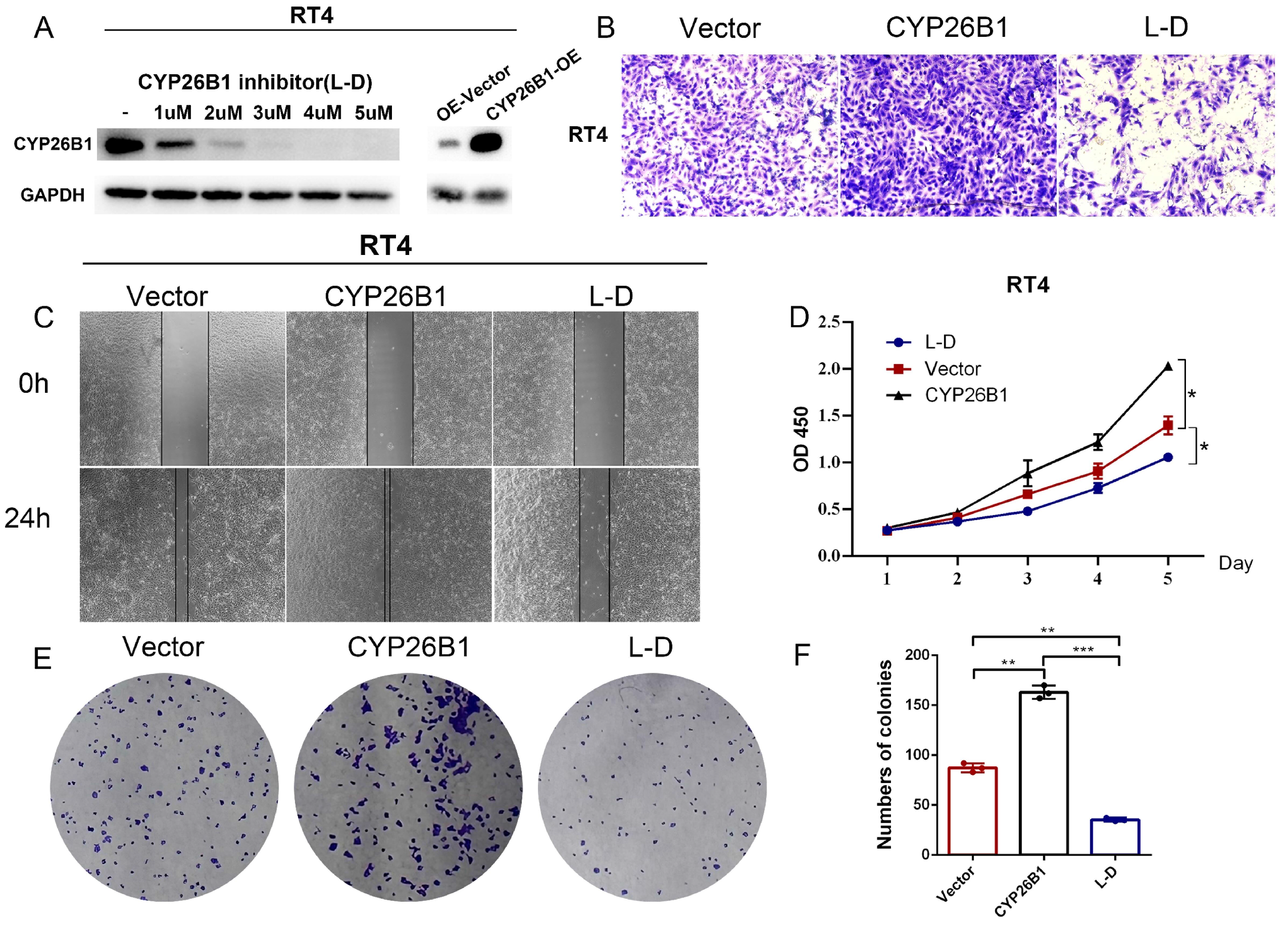
CYP26B1 promote BLCA migration and proliferation in RT4 cells. (A) Validation of inhibition and overexpression efficiency by western blot. (B) Transwell assay to explore cell migration ability in RT4 cell after overexpression and inhibition of CYP26B1. (C) Wound-healing assay to explore cell migration ability in RT4 cell after overexpression and inhibition of CYP26B1. (D) CCK8 assay to explore proliferation ability in RT4 cell after overexpression and inhibition of CYP26B1. (E-F) Colony formation assay to explore proliferation ability in RT4 cell after overexpression and inhibition of CYP26B1.

### Exploring the association between CRGs and the tumor microenvironment

The tumor microenvironment has become an important subject because of immunotherapy. Here, we explored the association between CRGs and the tumor microenvironment. CRGs were negatively correlated with tumor-infiltrating CD4^+^ T cells, Natural Killer T cells (NKT), and follicular helper T cells (Tfh) (**Figure 7A**) and positively correlated with neutrophils and Tregs. Thus, cuproptosis may be associated with the dysfunction of anti-tumor immunity. Furthermore, we evaluated the prognostic significance of CRGs in the IMvigor210 cohort to assess the potential association between immunotherapy response and candidate biomarkers. Univariate regression analysis (**Figure 7B**) indicated that *DLST* (*p*<0.01, HR=1.791) was significantly associated with shorter OS, whereas *LIPT1* (*p*<0.05, HR=0.896) was significantly associated with longer OS. Higher *DLST* expression levels served as a risk factor in patients treated with atezolizumab (**Figure 7C**). Thus, *DLST* may be associated with a dysfunctional or deserted immune microenvironment. *DLST* was positively correlated with various immune checkpoint molecules, *TGFB1* (correlation=0.273, *p*<0.0001), and *CD274* (correlation=0.11, *p*<0.05) in BLCA (**Figure 7D–F**). *DLST* was negatively correlated with immune stimulators, including *ICOSLG* (correlation=-0.298, *p*<0.0001) and *TNFRSF13B* (correlation=-0.218, *p*<0.0001) in BLCA (**Figure 7G–I**).

**Figure 7.**
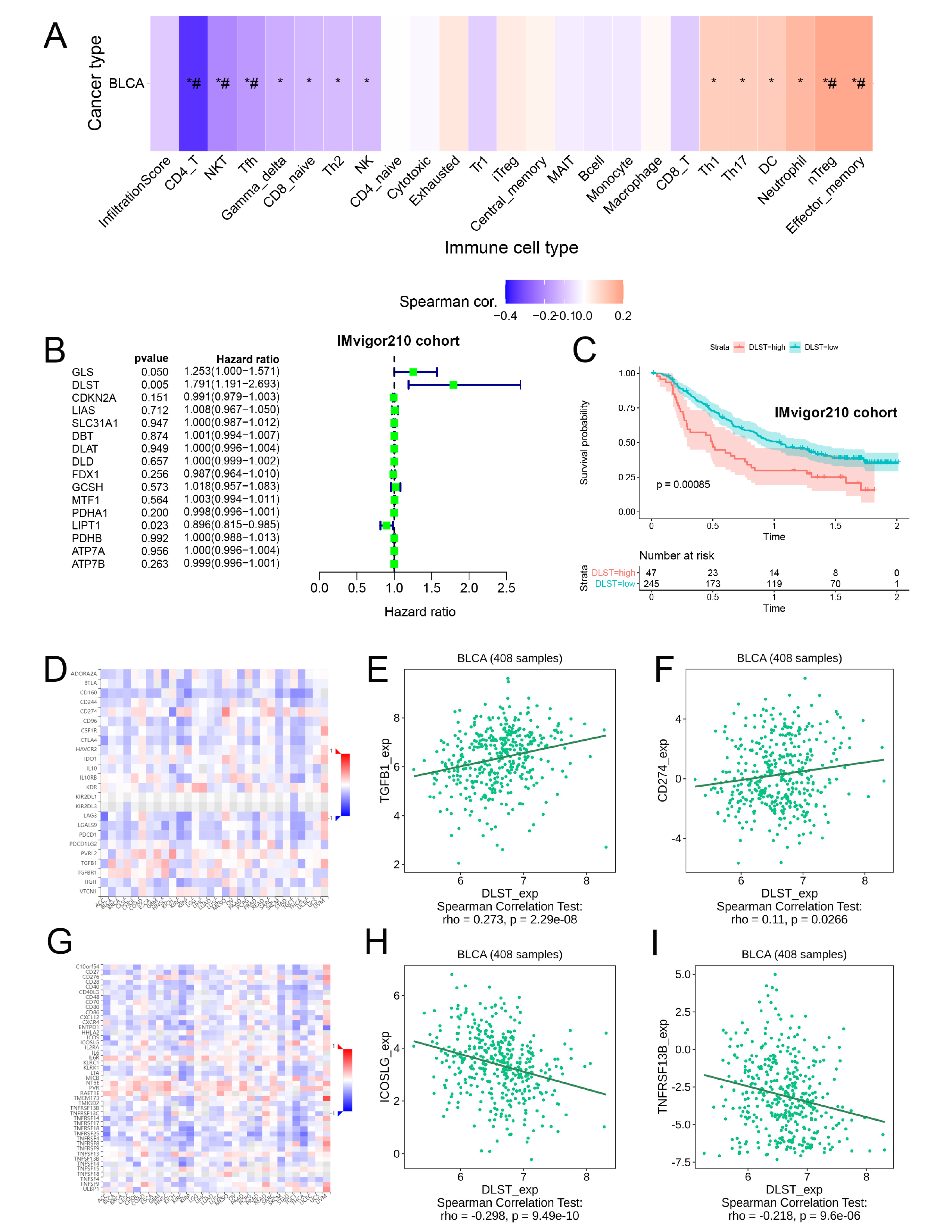
Exploring the association between CRGs and tumor microenvironment. (A) The association between CRGs and tumor microenvironment. (B) Univariate regression analysis to evaluate the prognostic significance of the CRGs in IMvigor210 cohort. (C) Overall survival curves of patients with high and low expression of DLST using the Kaplan-Meier method. (D) The heatmap showing the relationship between the expression of DLST and the immune checkpoints. (E and F) The relationship between the expression of DLST and the immune checkpoints TGFB1 and CD274, respectively. (G) The heatmap showing the relationship between the expression of DLST and the immune stimulators. (H and I) The relationship between the expression of DLST and the immune stimulators ICOSLG and TNFRSF13B.

### mIHC indicates a deserted immune microenvironment associated with DLST

Using the single-cell RNA dataset GSE130001, we explored the distribution of *DLST* in the BLCA tumor microenvironment. Most *DLST* expressions were localized in the epithelial cells (**Figure 8A–B**). **Figures 8C** and **8D** are representative images of the tumor microenvironment in the low and high *DLST* expression groups, respectively. The *DLST* expression levels were significantly higher in tumor tissues than in normal tissues (**Figure 8E**). Quantification of CD3, CD4, and CD8 signals indicated a significantly lower infiltration of CD3^+^, CD4^+^, and CD8^+^ T cells in the high *DLST* expression group.

**Figure 8.**
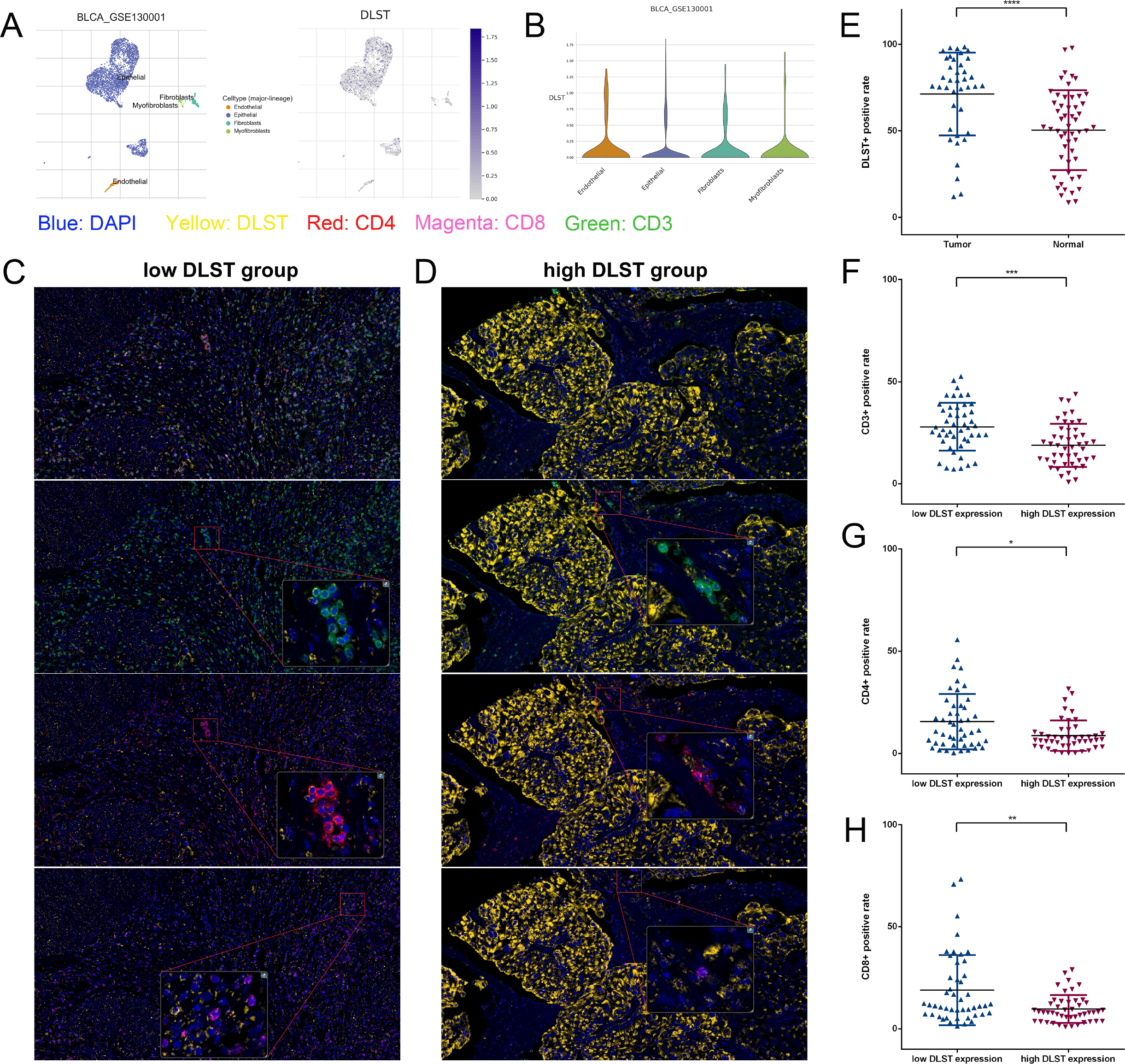
Multiple immunohistochemistry indicates a deserted immune microenvironment associated with DLS*T*. (A and B) The distribution of *DLST* in BLCA tumor microenvironment using single cell RNA dataset GSE130001. (C and D) Multiplex immunohistochemistry showing the expression level of DLST in bladder cancer tissues and adjacent normal tissues, respectively. (E and F) Multiplex immunohistochemistry showing the tumor infiltrating immune cells (including CD3^+^, CD4^+^ and CD8^+^ T cells) in tissues with high and low expression level of DLST, respectively. The scale bar of figures on the left is 200μm, while the scale bar of figures on the right is 50μm.

## Discussion

Cuproptosis, a novel cell death pathway proposed by Tsvetkov [4], is the direct binding of copper ions to the fatty acylated components of the TCA cycle, leading to abnormal aggregation of fatty acylated proteins and loss of iron-sulfur proteins. This further leads to a proteotoxic stress response and eventually cell death. Sixteen CRGs have been identified [4]. In this study, we explored the relationship between cuproptosis and BLCA. We downloaded the mutation and expression levels of CRGs in patients with BLCA from the cbioportal and TCGA databases, respectively, explored the potential function of CRGs in cancer using GSCA, and found that CRGs were closely related to the tumor cell cycle and apoptotic pathway. In addition, we evaluated the cuproptosis score of TCGA BLCA samples by LASSO Cox regression and divided the samples into two groups: high and low cuproptosis scores. Patients in the high cuproptosis score group had shorter OS, indicating that the established cuproptosis score model had an accurate prognostic ability.

We explored the relationship between CRGs and chemotherapy drug resistance in BLCA by analyzing the genetic differences in the high and low cuproptosis score group samples and the BLCA drug resistance-related dataset from the GEO database. We selected the intersection of the different genes from both groups and obtained 74 CRGs associated with chemoresistance. Finally, we screened the most significant DEG, *CYP26B1*. Cytochrome P450 family 26 (CYP26) belongs to the cytochrome P450 protein superfamily, which regulates the concentration of retinoic acid by inactivating all-trans retinoic acid into its hydroxylated form as a hydroxylase [23]. There are three enzymes in the mammalian CYP26 family: CYP26 subfamily A member 1 (CYP26A1), CYP26 subfamily B member 1 (CYP26B1), and CYP26 subfamily C member 1 (CYP26C1) [24]. *CYP26B1* is a susceptibility gene for esophageal squamous cell carcinoma [25] and increases the risk of oral squamous cell carcinoma [26]. However, its association with BLCA was unclear. High *CYP26B1* expression was associated with shorter OS and DSS in patients with BLCA, as well as with higher HRD and MSI scores, suggesting that *CYP26B1* is associated with platinum-based chemotherapy drugs and PARP inhibitor sensitivity and that its chemotherapy drug resistance is due to DNA mismatch repair. In addition, Transwell, wound healing, and CCK8 assays showed that *CYP26B1* was highly correlated with the proliferation and migration of BLCA cells. The above analysis revealed that the CRG *CYP26B1* is a key molecule in BLCA chemoresistance and may be an important target for its treatment.

We further explored the role of cuproptosis in the BLCA immune microenvironment. We first explored the correlation between CRGs, immune cells, and immune components and performed a prognostic analysis of CRGs using the BLCA immunotherapy cohort IMvigor210. *DLST* had the most significant prognostic effect, and patients with BLCA and high *DLST* expression had shorter OS, suggesting that *DLST* is associated with poor immunotherapy outcomes. DLST is a TCA cycle transferase in the α-ketoglutarate dehydrogenase complex (KGDHC) that catalyzes the conversion of α-KG to succinyl coenzyme A for energy production and macromolecule synthesis [27]. DLST is an important mediator of T-cell acute lymphoblastic leukemia (T-ALL) and triple-negative breast cancer [28–30] and promotes the aggressiveness of neuroblastoma [31]. However, the role of DLST in the tumor immune microenvironment and BLCA has not yet been reported. In our study, mIHC indicated that the expression level of DLST was relatively low in adjacent normal tissues, whereas significantly elevated in tumor tissues. High DLST expression levels were associated with low immune infiltration. In contrast, many tumor-infiltrating immune cells, including CD3^+^, CD4^+^, and CD8^+^ T cells, were present in tumors with low DLST expression. Thus, high DLST expression may imply an immunosuppressive microenvironment and patients with high DLST expression may not benefit from immunotherapy.

## Conclusions

In this study, we explored the relationship between cuproptosis, a novel cell death pathway, and BLCA. We first investigated the variation and expression levels of CRGs in patients with BLCA and used LASSO Cox regression to construct a BLCA cuproptosis score model for predicting the prognosis of patients with BLCA. We then used the BLCA chemoresistance dataset in the GEO database to identify CYP26B1, a key cuproptosis molecule that promotes chemoresistance in BLCA. Subsequent experiments and bioinformatics analysis confirmed the crucial role of CYP26B1 in BLCA chemoresistance and the malignant phenotype. In addition, we used a BLCA immunotherapy cohort to screen for DLST, a molecule closely associated with the immune microenvironment of BLCA and confirmed that bladder cancer tissues with high expression of DLST are associated with low immune cell infiltration through mIHC. Our findings provide possible reasons for the poor immunotherapeutic outcome in patients with BLCA and provide potential targets for immunotherapy.

## Declarations

### Acknowledgments

We thank TCGA and GEO databases for providing NGS data and clinical information on BLCA.

### Author Contributions

The work introduced here was performed jointly by all authors. Cong-hui Han determined the research idea, revised the draft and managed the whole study. Zhen-duo Shi, Kun Pang, and Lin Hao analyzed and interpreted the data for the work. Wei Wang helped with experimental validation and data collection. Jun Gu and Lijun pang drafted the manuscript, analyzed the data, constructed the model, carried out related experiments, and explained the results.

### Data Availability Statement

Somatic variation data of the TCGA-BLCA cohort were obtained from the cBioPortal for Cancer Genomics (http://cbioportal.org). Gene expression profiles and corresponding clinical information of 411 BLCA tumor samples were obtained from TCGA database (https://portal.gdc.cancer.gov/). DEGs between chemotherapy-sensitive and chemotherapy-resistant mouse bladder cancer and dataset GSE192575 were obtained from https://www.ncbi.nlm.nih.gov/geo/query/acc.cgi?acc=GSE192575. We also obtained the IMvigor210 cohort from http://research-pub.gene.com/IMvigor210CoreBiologies, which contains locally advanced and metastatic uroepithelial cancer patients treated with atezolizumab. Single-cell RNA-seq profiling of two bladder cancer specimens was performed using the GSE130001 dataset (https://www.ncbi.nlm.nih.gov/geo/query/acc.cgi?acc=GSE130001).

### Ethics Statement

Ethics approval and consent to participate in the current study were obtained from the ethics committee of Suzhou Medical College of Soochow University. All the patients enrolled for analysis in this research had signed consent forms before they underwent surgical treatment, which allows us to use their samples for scientific research. Ethics committee of Suzhou Medical College of Soochow University will also perform an ethical review before researches begin and the collection of the patients’ samples is following general principles of the WMA Declaration of Helsinki.

### Consent for publication

Not applicable.

